# Identification and Rescue of Congenital Hyperinsulinism-Associated *ABCC8* Mutations that Impair K_ATP_ Channel Trafficking

**DOI:** 10.1101/2025.05.18.654760

**Authors:** Assmaa ElSheikh, Yi-Ying Kuo, Kara E. Boodhansingh, Zhongying Yang, Charles A. Stanley, Diva D. De Leon, Show-Ling Shyng

## Abstract

ATP-sensitive potassium (K_ATP_) channels composed of Kir6.2 and sulfonylurea receptor 1 (SUR1) couple glucose metabolism with insulin secretion in pancreatic β-cells and are vital to glucose homeostasis. Loss-of-function mutations in SUR1 and Kir6.2, encoded by *ABCC8* and *KCNJ11*, respectively are the commonest causes of severe persistent hypoglycemia in infants and children seen in the rare disease congenital hyperinsulinism (HI). The N-terminal transmembrane domain, TMD0, and the linker immediately C-terminal to TMD0, L0, of SUR1 (TMD0/L0) forms direct contact with Kir6.2 in K_ATP_ channels. Mutations in SUR1-TMD0 often impair K_ATP_ channel trafficking to the plasma membrane, causing severe disease unresponsive to treatment by the K_ATP_ activator diazoxide; however, surface expression and function of many such mutant channels can be rescued by reversible K_ATP_ inhibitor pharmacochaperones. Here, we identified seven new SUR1 missense mutations in TMD0/L0 from HI patients unresponsive to diazoxide and investigated their effects on K_ATP_ channel expression, function, and response to pharmacochaperones. All seven mutations, N32K, Y124F, P133R, W143R, L171P, G228D, and Y230C, reduced channel function in Rb^+^ efflux assays. Further characterization by immunoblotting, immunostaining and electrophysiology revealed that Y124F primarily causes defective channel gating, while the others impair channel trafficking to different extents. The trafficking mutations showed varied response to surface expression and function rescue by the reversible K_ATP_ inhibitor pharmacochaperones, tolbutamide and Aekatperone. The study underscores the critical role of SUR1-TMD0/L0 in K_ATP_ expression and gating. It further highlights the importance of detailed biochemical and functional studies of mutant channels in understanding their pathogenic roles and response to potential pharmacological therapies.

## Introduction

In pancreatic β-cells, ATP-sensitive potassium (K_ATP_) channels formed by four pore-forming Kir6.2 subunits and four regulatory sulfonylurea receptor 1 (SUR1) subunits regulate insulin secretion by coupling changes in intracellular ATP/ADP ratios from glucose metabolism with membrane potential (1–3). When blood glucose levels are low, the relatively low intracellular ATP/ADP ratio allows K_ATP_ channels to open, which keeps the β-cell membrane in a hyperpolarized state to prevent insulin secretion. When blood glucose levels are elevated, the intracellular ATP/ADP ratio increases to close K_ATP_ channels, leading to membrane depolarization, opening of voltage-gated Ca^2+^ channels, and insulin release. This finely tuned process ensures insulin secretion is appropriately regulated by glucose to maintain euglycemia.

Congenital hyperinsulinism (HI) is a rare disease characterized by persistent insulin secretion despite life-threatening hypoglycemia (4, 5). Variants in many genes involved in fuel metabolism and the insulin secretory pathway in pancreatic β-cells have been linked to HI (6). However, the most frequent causes of the disease are loss-of-function mutations in *KCNJ11* and *ABCC8*, which respectively encode the K^+^-conducting pore subunit Kir6.2 and the regulatory sulfonylurea receptor 1 (SUR1) subunit of the β-cell K_ATP_ channel (referred to as K_ATP_-HI) (4, 6, 7). Loss of K_ATP_ channel function renders persistent β-cell depolarization and insulin secretion even when blood glucose is dangerously low (3). Prompt diagnosis and treatment based on comprehensive biochemical and functional evaluation of new K_ATP_ variants identified in HI is critical for designing better therapeutic approaches to prevent severe, long-term complications of the disease.

Numerous HI-associated genetic alterations in the β-cell K_ATP_ channel have been identified, mostly in the large SUR1 subunit encoded by *ABCC8* (8, 9). A fraction of these have been evaluated for their impact on the biochemical and biophysical properties of K_ATP_ channels and verified as pathogenic. These studies show that pathogenic mutations can impair channel gating, channel trafficking to the cell surface, or both (7, 10–13). Most gating mutations reduce or abolish channel response to MgADP (7, 14), which is the physiological K_ATP_ activator at low glucose (15) by binding to SUR1 to stimulate Kir6.2 opening (16). Patients with gating mutations can sometimes be successfully treated with the K_ATP_ channel opener diazoxide, the only FDA-approved drug for HI (4, 7). However, patients with trafficking mutations that greatly reduce K_ATP_ surface expression typically present severe disease that is unresponsive to diazoxide, often requiring pancreatectomy to curb hypoglycemia (4, 7). For these patients, alternative therapeutic strategies that address the underlying molecular defects are needed.

Mutations that impair K_ATP_ trafficking are found throughout SUR1 and Kir6.2; however, a great number of them are located in the N-terminal transmembrane domain of SUR1, called TMD0, and the downstream cytoplasmic linker, called L0 (17). The TMD0-L0 domain of SUR1 is the primary assembly domain with Kir6.2 as revealed by cryoEM structures of the K_ATP_ channel (16, 18). We have previously shown that trafficking defects caused by mutations in TMD0 of SUR1 may be corrected by small molecules known to inhibit K_ATP_ channels, which we refer to as K_ATP_ pharmacochaperones (17). However, the inhibitory pharmacochaperones need to be removed from rescued surface mutant channels to recover channel function for clinical applications. K_ATP_ pharmacochaperones reported in early work either inhibit the channel irreversibly, such as the high affinity sulfonylurea glibenclamide (GBC) and the glinide repaglinide (19, 20), or have other known off-targets, such as the low affinity sulfonylurea tolbutamide (Tolb) (19) and the anticonvulsant carbamazepine (21). This prompted us to recently undertake a structure-based drug discovery effort to search for additional reversible K_ATP_ inhibitors, which led to the identification of a novel reversible K_ATP_ inhibitor pharmacochaperone named Aekatperone (AKP) (22). AKP was shown to rescue surface expression of several previously identified HI-causing SUR1-TMD0 trafficking mutants and be easily washed out to recover the function of rescued mutant channels (22), making it a promising potential therapeutic for HI caused by diazoxide-unresponsive K_ATP_ trafficking mutations.

In this study, we investigated the effects of previously unreported or uncharacterized SUR1 mutations, the majority in TMD0-L0, identified from diazoxide-unresponsive HI patients on the expression and function of recombinant channels expressed in COSm6 cells. Moreover, we evaluated the effects of the newly identified K_ATP_ pharmacochaperone AKP on those variants that showed impaired trafficking to the plasma membrane. The study expands our knowledge on the correlation between genotype and clinical, molecular phenotype in HI with SUR1 mutations and those that are candidates for potential future pharmacological chaperone therapy.

## Results

### Identification of HI-associated ABCC8 variants

*ABCC8* single-nucleotide variants were identified in seven diazoxide-unresponsive CHI patients. These include seven previously unreported mutations that result in missense mutations in TMD0-L0 of SUR1: N32K, Y124F, P133R, W143R, L171P, G228D, and Y230C (Figure 1, Table 1).

**Figure 1.**
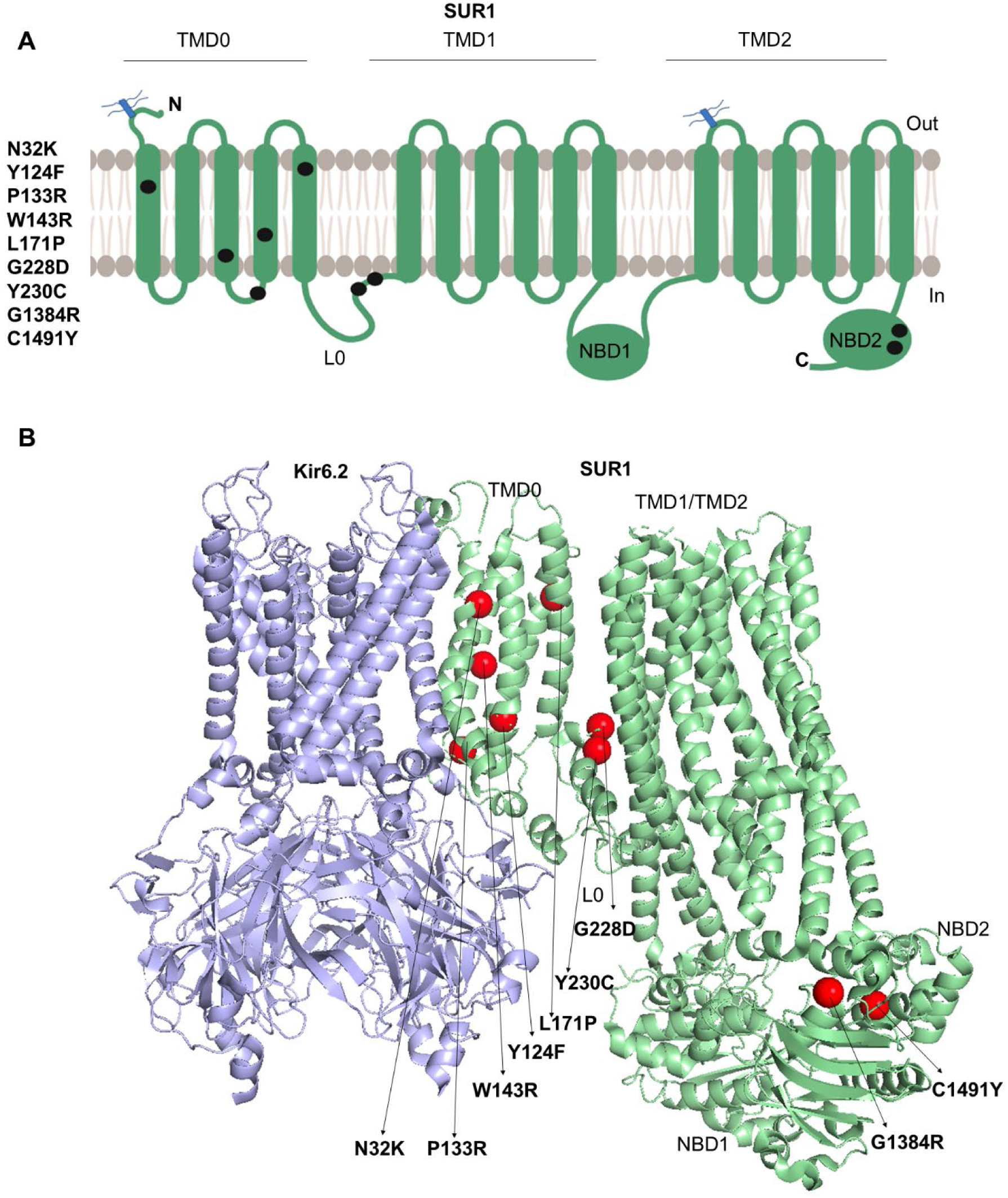
Positions of the newly identified Disease-Causing K_ATP_ Channel Mutations. **(A)** The positions of the seven SUR1 mutations analyzed in this study are indicated on a SUR1 topology model (49). (B) The mutation sites are shown in the K_ATP_ channel cryo-EM structure (PDB ID 6BAA) as red spheres within the TMD0/L0 and NBD2 domains (mutations in NBD2 are referred to as non-TMD0/L0 mutations here in the paper) of the SUR1 subunit. For clarity, only one SUR1and four Kir6.2 subunits are shown. SUR1 is colored in light green and Kir6.2 in blue.

**Table 1.** Genetic and clinical data of patients bearing the TMD0 mutations in *ABCC8* gene.

| Nucleotide | Change | Codon | Amino acid change | Associated mutations | Histology | Parent of origin |
| --- | --- | --- | --- | --- | --- | --- |
| 96 | C → G | 32 | N → K | *ABCC8*: Q954X (maternal) | Diffuse | Paternal |
| 371 | A → T | 124 | Y → F | *ABCC8*: C1491Y (maternal) | Diffuse | Non-maternal |
| 398 | C → G | 133 | P → R | none | Focal | Paternal |
| 427 | T → C | 143 | W → R | none | Focal | De novo |
| 512 | T → C | 171 | L → P | none | Focal | Paternal |
| 683 | G → A | 228 | G → D | *ABCC8*: c.3577delG (paternal) | Diffuse | Maternal |
| 689 | A → G | 230 | Y → C | *ABCC8*: G1384R (paternal) | Diffuse | Maternal |

Four of these: N32K, Y124F, G228D, and Y230C, were found in patients diagnosed with the diffuse form of HI who also carry a second mutant allele inherited from the opposite parent: Q954X, C1491Y, a frameshift mutation D1193Mfs resulting from c.3577delG, and G1384R, respectively (Table 1) (9). Q954X and D1193Mfs have been reported before (9) and are predicted to result in truncated or frameshifted non-functional SUR1. G1384R has also been reported previously but its impact on K_ATP_ channels has not been characterized (9), and C1491Y is a new mutation; both are in the second nucleotide binding domain (NBD2) of SUR1 (Figure 1). The other three TMD0 mutations: P133R, W143R, and L171P, were associated with focal HI, and were recessively inherited from the paternal parent or arose *de novo*. Currently, there are no reliable *in silico* methods to accurately predict the clinical significance of new *ABCC8* variants in HI. The problem becomes even more complex when they occur in conjunction with a second mutant allele. As such, direct experimental evidence is needed to understand the functional impact of genetic variants on K_ATP_ channel and the underlying molecular mechanisms.

### Reduced function of K_ATP_ channels containing the newly identified SUR1 missense mutations

To directly elucidate the effects of the newly identified SUR1 mutations on K_ATP_ channel function, we performed biochemical and functional analyses using recombinant channels transiently expressed in COSm6 cells, which do not express endogenous K_ATP_ channels, thus providing a clean background to study mutant channels.

First, we assessed the pathogenic roles of the mutations in HI by Rb^+^ efflux assays, which use Rb^+^ as a surrogate for K^+^ to report K_ATP_ channel activity in intact cells (23). COSm6 cells were co-transfected with human WT or mutant SUR1 and WT Kir6.2 cDNAs, preloaded with Rb⁺, and subjected to Rb⁺ efflux measurement over a 30-minutes period in Ringer’s solution that contained metabolic inhibitors (2.5 μg/ml oligomycin and 1 mM 2-deoxy-D-glucose; referred to as MI hereinafter) which mimic the effect of hypoglycemia to decrease intracellular ATP/ADP ratios, thereby activating K_ATP_ channels (see **Experimental procedures**). To evaluate mutant channel response to diazoxide, a K_ATP_ channel opener and the frontline medical treatment for HI, Rb^+^ efflux assays were also performed in Ringer’s solution containing 0.2 mM diazoxide. All seven TMD0/L0 mutations, N32K, Y124F, P133R, W143R, L171P, G228D, and Y230C, significantly reduced Rb^+^ efflux in response to MI or diazoxide compared to WT but to varying degrees (Figure 2). Among them, N32K, W143R, L171P, and G228D showed nearly no stimulation by MI or diazoxide. The P133R mutation showed residual response to MI (∼50% that of WT fractional efflux) but little stimulation by diazoxide. In contrast, Y124F and Y230C had significant partial response to both MI (∼70% that of WT) and diazoxide (40% and 60% of WT, respectively).

**Figure 2.**
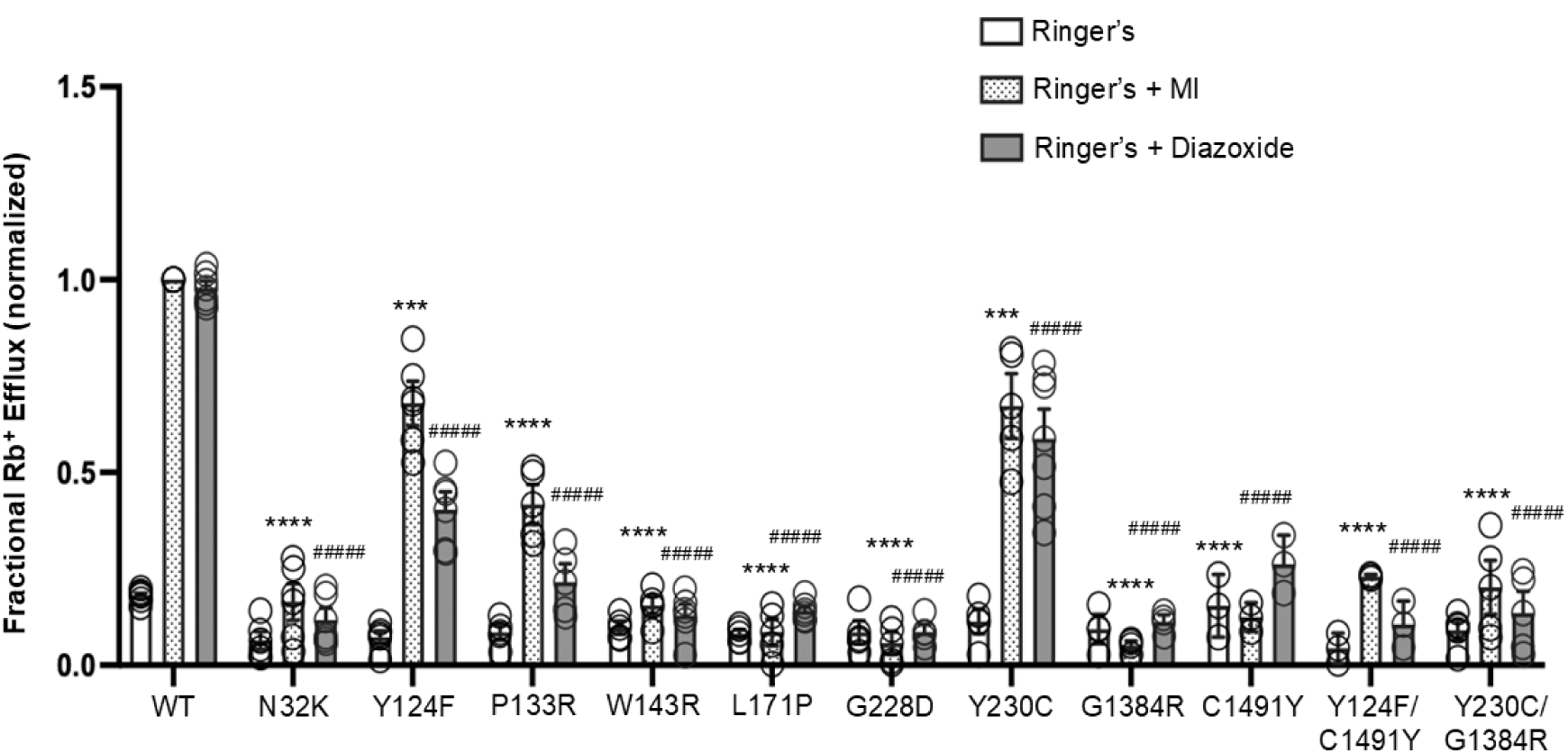
Functional Characterization of Pathogenic K_ATP_ Channel Mutations. Rb^+^ efflux assay results for COSm6 cells expressing WT and mutant K_ATP_ channels. To model the heterozygous state of Y230C/G1384R or Y124F/C1491Y, the two human SUR1 mutant constructs were co-expressed at a 1:1 ratio (see **Experimental procedures**). Fractional Rb⁺ efflux was measured under baseline conditions (Ringer’s solution), metabolic inhibition, (2.5 μg/ml oligomycin and 1 mM 2-deoxy-D-glucose) (Ringer’s + MI), or in the presence of 200 μM diazoxide (Ringer’s + Diazoxide). Data were normalized to the WT response under MI conditions after subtracting background Rb⁺ efflux from untransfected (UT) cells. Each bar represents the mean ± SEM from at least three biological replicates, with individual data points shown as open circles. Statistical significance was determined at α = 0.05 using one-way ANOVA followed by Dunnett’s multiple comparisons test. Comparisons to WT under MI conditions are indicated as follows: ****p < 0.0001, ***p < 0.001, **p < 0.01, *p < 0.05; comparisons to the WT response under diazoxide treatment are denoted as ####p < 0.0001.

The discordance between the mild effects of Y124F and Y230C and the severe disease/diazoxide unresponsiveness observed in patients may be attributable to the presence of a second mutation, C1491Y and G1384R, respectively, on the opposite SUR1 allele. We therefore examined the impact of the two non-TMD0/L0 mutations on channel response to MI and diazoxide, both alone, and in combination with Y124F for C1491Y and Y230C for G1384R. Rb^+^ efflux assays showed that indeed C1491Y and G1384R rendered channels unresponsive to MI and diazoxide. Interestingly, in cells co-expressing Y124F-SUR1 and C1491Y-SUR1 at 1:1 cDNA ratio, to model heterozygous expression in vitro, the efflux levels in MI or diazoxide were below 50% of those seen in cells expressing Y124F alone, suggesting the C1491Y mutation has a dominant-negative effect over the less severe Y124F mutation (Figure 2). Likewise, we observed a dominant-negative effect of G1384R over Y230C in cells co-expressing both mutant SUR1 proteins at 1:1 ratio (Figure 2). These results indicate that in patients with the Y124F/C1491Y and the Y230C/G1384R compound mutations, C1491Y and G1384R have a dominant role in dictating the severe, diazoxide-unresponsive clinical phenotype.

### Effects of the mutations on K_ATP_ Channel maturation

The above mutations could reduce K_ATP_ function by impairing channel biogenesis and trafficking to the cell surface, causing abnormal gating response to intracellular ATP and ADP, or both. To evaluate effects of the mutations on K_ATP_ channel biogenesis, western blot analysis of SUR1 was performed. SUR1 contains two N-linked glycosylation sites, which undergo core glycosylation within the endoplasmic reticulum (ER) (24, 25). Upon successful assembly with Kir6.2 to form octameric K_ATP_ channels and subsequent exit from the ER, SUR1’s N-linked glycosylation sites are further modified in the Golgi to give rise to complex-glycosylated SUR1 before trafficking to the cell surface. These core and complex glycosylated SUR1 can be separated by SDS-PAGE, appearing as a lower (immature) band and an upper (mature) band, respectively (26, 27). Since only fully assembled channels pass ER quality control, the relative abundance of the mature and immature bands can be used to approximate channel maturation efficiency. COSm6 cells co-transfected with WT SUR1 plus WT Kir6.2 cDNAs or mutant SUR1 plus WT Kir6.2 cDNAs were subjected to western blot analysis (Figure 3A). Of the seven SUR1 TMD0/L0 mutations, Y124F-SUR1 is the only one which exhibited a relatively normal upper band comparable to WT-SUR1; all the others, namely, N32K-, P133R-, W143R-, L171P-, G228D- and Y230C-SUR1 showed clearly reduced upper SUR1, indicating impaired channel biogenesis and trafficking. The relative upper band abundance as a percentage of the total band abundance (both upper plus lower) for each mutation across at least three independent experiments was quantified by densitometry using ImageJ. The analysis confirmed that except for Y124F (20 ± 13% reduction), all other mutations caused a significant reduction in the upper SUR1 band (95 ± 1%, 79 ± 5%, 95 ± 2%, 97 ± 1%, 98 ± 3%, and 60 ± 13% for N32K, P133R, W143R, L171P, G228D, and Y230C, respectively; Figure 3B).

**Figure 3.**
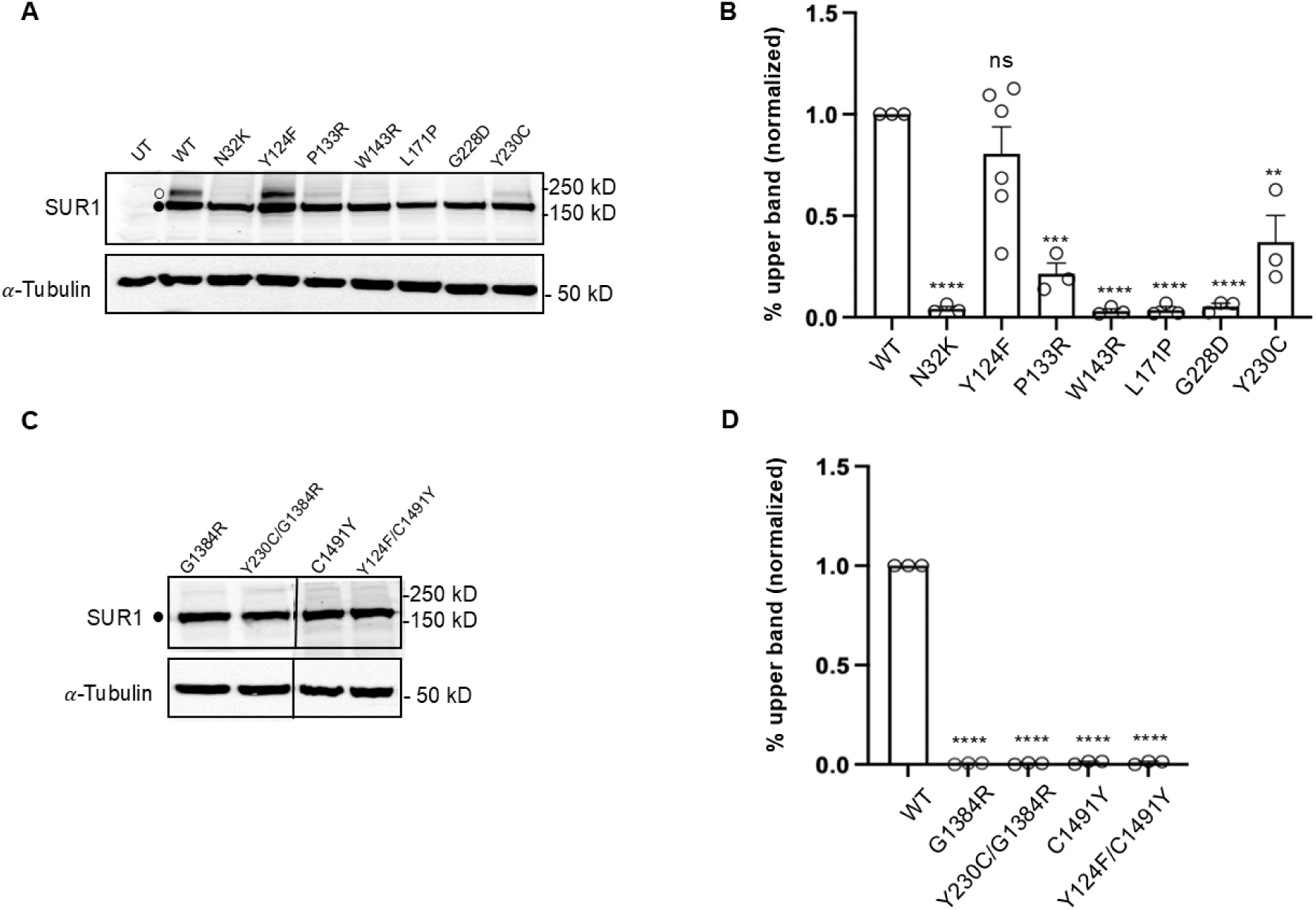
Biochemical Characterization of the Maturation Efficiency of the TMD0/L0 and non-TMD0/L0 SUR1 mutations. (A, C) Western blots of SUR1 from COSm6 cells co-transfected with human WT Kir6.2 and WT or mutant SUR1 cDNA. In (A), untransfected cells (UT) and cells expressing WT channels were included for comparison. The solid circle marks the core-glycosylated (immature) form of SUR1, and the open circle marks the complex-glycosylated (mature) form of SUR1. In (C), COSm6 cells co-transfected with human WT Kir6.2 and mutant SUR1 cDNAs (G1384R, Y230C/G1384R (1:1), C1491Y, or Y124F/C1491Y (1:1). In both panels, the tubulin blot below serves as the loading control. Molecular weight markers are shown to the right of the blots, with units in kD. (B, D) Bar graphs illustrate the quantification of the relative density of the upper band (%) in comparison to the total expression (combined density of upper and lower bands), as measured using ImageJ densitometry from at least three independent experiments. Each circle represents an independent experiment. The data were normalized to the percentage of the upper band observed in WT channels. Statistical significance was assessed at α = 0.05 using one-way ANOVA followed by Dunnett’s multiple comparisons test. Comparisons to WT are represented as follows: \*\*\*\**p* < 0.0001, \*\*\**p* < 0.001, \*\**p* < 0.01, and ns (not significant).

We also determined the maturation efficiency of the two non-TMD0/L0 SUR1 mutations, G1384R and C1491Y by western blots. Neither G1384R nor C1491Y had detectable upper mature SUR1 band (Figure 3C, D). When C1491Y was co-expressed with Y124F, no upper SUR1 band was observed (Figure 3C, D), suggesting a dominant-negative effect of C1491Y over Y124F on SUR1 maturation. Similarly, co-expression of G1384R with Y230C eliminated the weak upper SUR1 band seen in cells expressing Y230C alone (Figure 3C), consistent with a dominant-negative effect of G1384R on Y230C maturation. These results indicate that C1491Y and G1384R exert their functional dominant-negative effects over Y124D and Y230C by preventing channel maturation thereby surface expression.

### Y124F-SUR1 reduces K_ATP_ channel response to MgADP

Since the Y124F-SUR1 mutation showed no significant effect on SUR1 maturation in western blots (Figure 3A, B) but significantly reduced channel activity in Rb^+^ efflux assays (Figure 2), we hypothesized that the mutation reduces channel function by altering channel gating properties. Physiological activity of K_ATP_ channels is determined by the balance of channel inhibition by ATP via Kir6.2 and channel activation by MgADP via SUR1 (2, 16, 28). A reduced sensitivity to MgADP stimulation is the most common defect observed in HI-associated K_ATP_ channel mutations (14, 15). To test whether the Y124F mutation alters the channel’s sensitivity to MgADP, we compared channel activity in 0.1 mM MgATP alone to that in 0.1 mM MgATP plus 0.5 mM MgADP using inside-out patch-clamp recordings. As shown in Figure 4A, B, the Y124F mutation significantly reduces the channel’s response to MgADP stimulation compared to WT channels. Additionally, the mutation markedly decreases the channel’s response to diazoxide (Figure 4C, D). These results corroborate the Rb^+^ efflux results (Figure 2) and demonstrate that the Y124F mutation impairs channel gating response to the physiological activator MgADP and pharmacological opener diazoxide.

**Figure 4.**
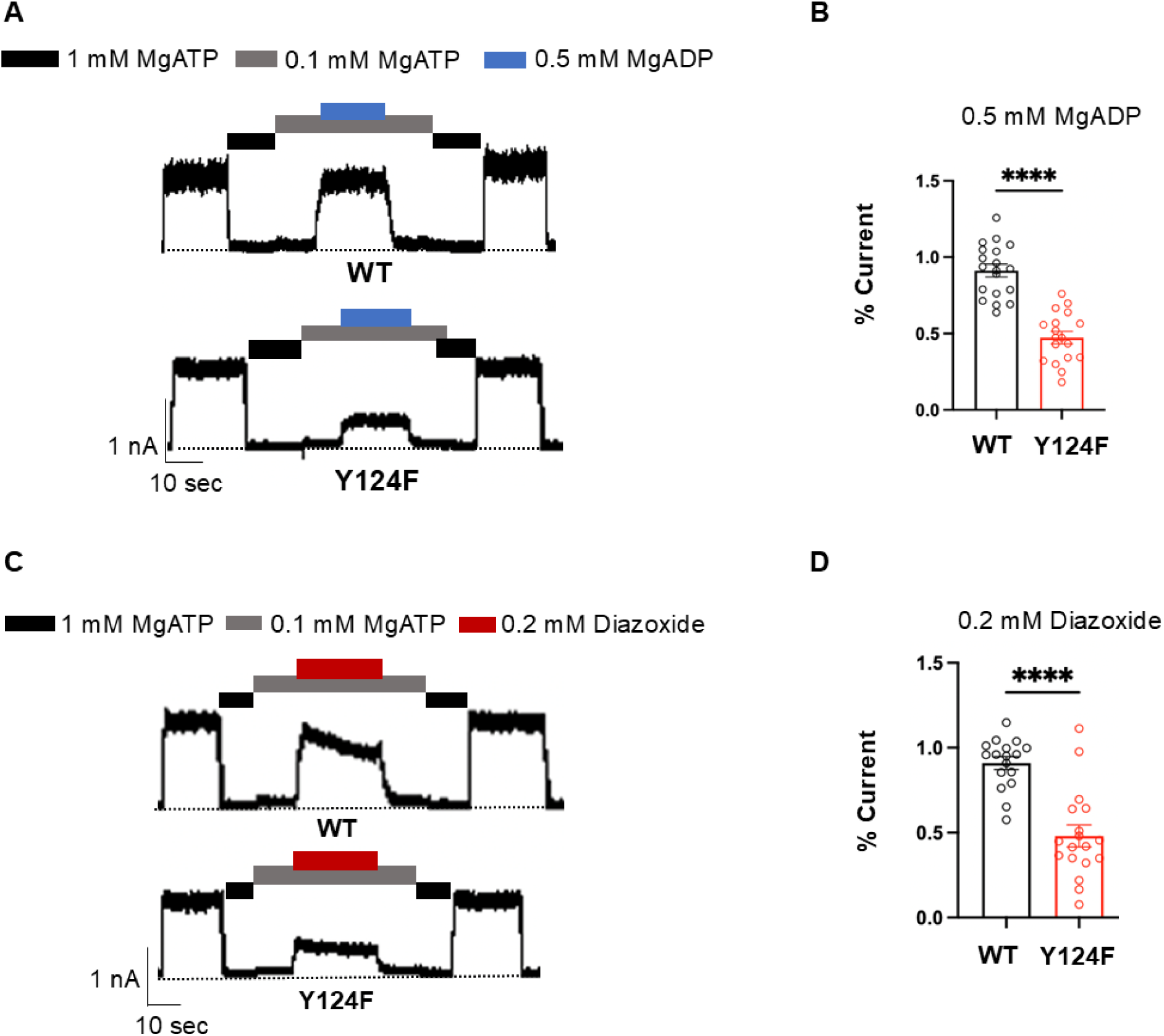
The Y124F SUR1 mutation is associated with a gating defect in K_ATP_ channels. (A, C) Representative inside-out patch-clamp recordings (holding potential of −50 mV; upward deflections indicate inward currents; black dashed lines denote the 0 current baseline in 1 mM MgATP) for human WT SUR1/Kir6.2 channels and Y124F SUR1/Kir6.2 channels. The channels were activated by 0.5 mM MgADP or 0.2 mM Diazoxide, respectively, as indicated by the bars above the recordings. (B, D) Bar graphs showing quantitative analysis of all recordings like those shown in (A) and (C), with significance assessed by *t*-test (α = 0.5). Significance levels indicated: \*\*\*\**p* < 0.0001.

### Pharmacological rescue of HI-causing K_ATP_ trafficking mutants

K_ATP_ channel inhibitors, including sulfonylureas and glinides widely used to treat type 2 diabetes, are known to act as pharmacochaperones to correct K_ATP_ trafficking defects caused by mutations in the SUR1-TMD0 domain (17, 19, 29, 30). CryoEM structures have elucidated that these compounds promote K_ATP_ channel trafficking to the plasma membrane by facilitating SUR1 and Kir6.2 assembly. The structural knowledge has led to our recent identification of a novel reversible K_ATP_ inhibitor, Aekatperone (AKP), which we have shown to rescue several previously reported SUR1-TMD0 trafficking mutants and be easily removed from rescued surface channels to recover their function (22). We asked whether the newly identified HI-associated TMD0/L0 mutations here: N32K, P133R, W143R, L171P, G228D, and Y230C (Figure 3A, B), could also be rescued by AKP.

Western blot analysis was used to assess whether treatment with AKP could improve the maturation of mutant SUR1. For comparison, we also included two other known K_ATP_ pharmacochaperones: the high-affinity sulfonylurea GBC and the low-affinity sulfonylurea tolbutamide, Tolb. COSm6 cells transiently expressing mutant channel proteins were treated overnight (16 h) with vehicle control (0.1% DMSO), 10 μM GBC, 200 μM Tolb, or 100 μM AKP. The concentrations were chosen to give maximum effects based on prior studies (19, 22). WT SUR1/Kir6.2 was included as a positive control in each blot. The results, presented in Figure 4A, show that while all six SUR1 mutants had no or faint mature upper band in the vehicle (0.1% DMSO) treated control, drug treatment led to the appearance or increased the upper SUR1 band in some of the mutants. Specifically, the N32K, P133R, W143R, and Y230C mutations showed clear albeit variable response to the three pharmacochaperones (Figure 5A), while the L171P and G228D mutations showed no obvious response to any of the three pharmacochaperones.

**Figure 5.**
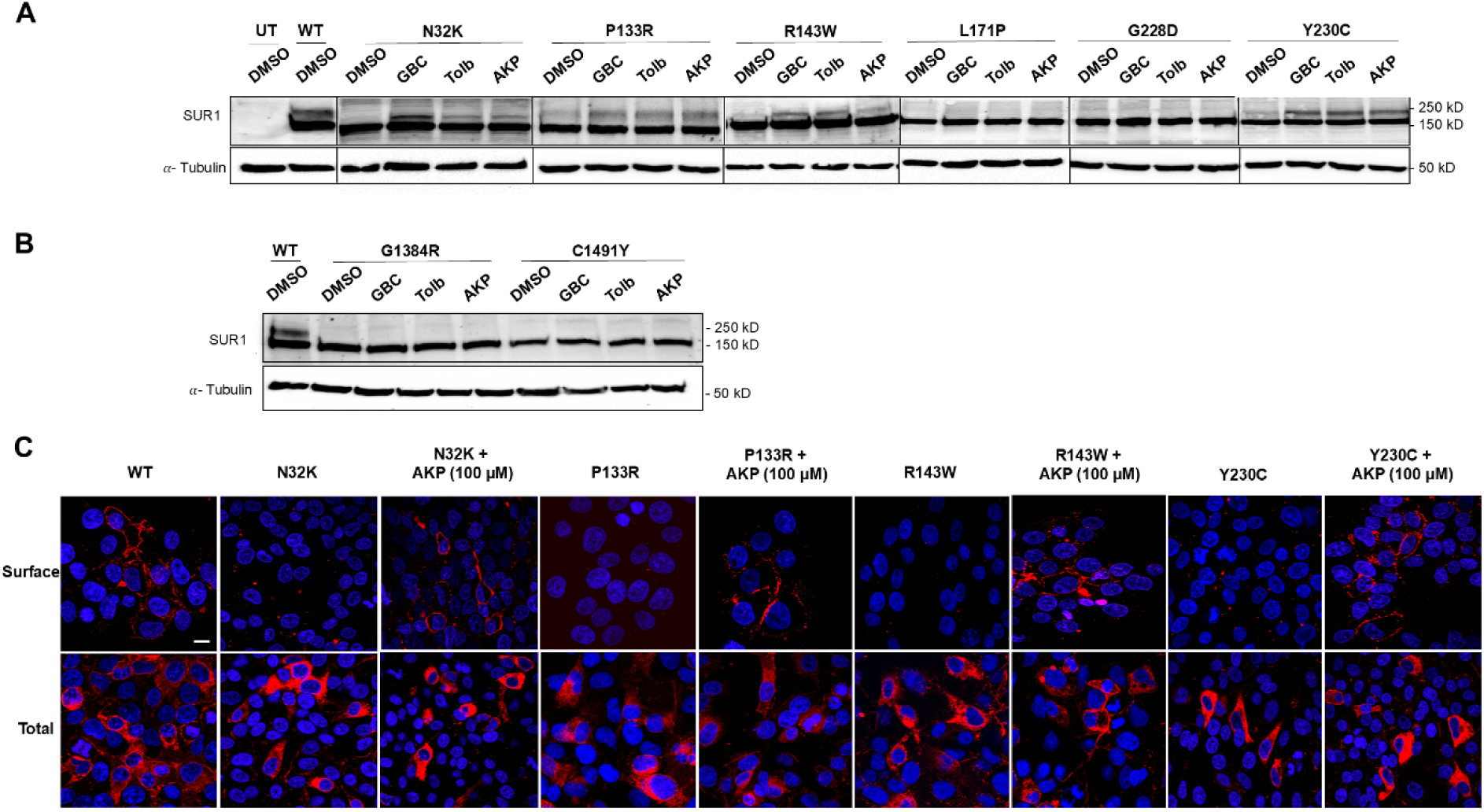
Pharmacological correction of SUR1 processing defects caused by the newly identified SUR1 mutations. (A, B) Western blots of SUR1 from COSm6 cells co-transfected with WT Kir6.2 and mutant SUR1 cDNA, treated with 0.1% DMSO (vehicle control), 10 μM Glibenclamide (GBC), 200 μM Tolbutamide (Tolb), or 100 μM Aekatperone (AKP) for 16 hours. TMD0/L0 mutations are shown in (A) and non-TMD0/L0 mutations are shown in (B). Untransfected cells (UT) and cells expressing WT channels were included as controls and only shown in (A). Tubulin blots below served as loading controls. The thin lines demarcate different sections of the blots or different blots. Molecular weight markers are shown on the right (in kD). (C) Top panels: Surface expression of human constructs of FLAG-tagged SUR1 (f-SUR1) WT or N32K, P133R, W143R, or Y230C trafficking mutants in COSm6 cells co-transfected with WT human Kir6.2. Cells expressing trafficking mutants were treated with 100 μM Aekatperone (AKP) for 16 hours. Surface localization was determined by immunostaining the extracellular FLAG epitope (red) in non-permeabilized cells. Nuclei were counterstained with DAPI (blue). Bottom panels: Total cellular expression of FLAG-tagged SUR1. Cells were fixed and permeabilized before incubation with primary and secondary antibodies. Images represent stacked sections from Olympus Fluoview confocal microscopy, with DAPI counterstaining. Scale bar, 15 μm. WT channels were used as controls for comparison.

Previous studies found that trafficking mutations located in the SUR1-ABC core outside TMD0/L0 do not respond to pharmacochaperone rescue (17, 19, 30). Consistent with this pattern, we do not see any increase in upper SUR1 band in cells expressing G1384R or C1491Y treated with GBC, Tolb, or AKP (Figure 5B).

To validate that the increased mature SUR1 band observed in N32K, P133R, W143R, and Y230C following AKP treatment corresponds to enhanced surface expression of the mutant K_ATP_ channels, we performed immunofluorescent staining to directly detect channel surface expression. COSm6 cells were co-transfected with Kir6.2 and N-terminally (extracellular) FLAG-tagged SUR1 (31). Surface K_ATP_ channels were detected by incubating live cells in a solution containing an anti-FLAG antibody at 4°C (see **Experimental procedures**), a temperature that halts membrane trafficking. As expected, cells transfected with the WT K_ATP_ channel constructs exhibited strong surface staining. In contrast, cells transfected with the N32K, P133R, W143R, or Y230C mutant constructs and treated with DMSO displayed minimum surface staining. However, staining of fixed and permeabilized cells revealed abundant intracellular fluorescence localized to the perinuclear region, consistent with ER retention of the mutant channels (Figure 5C, lower panels). Upon treatment with AKP (100 μM) for 16h, there was a marked increase in surface staining for all four mutants (Figure 5C, top panels), providing direct evidence that AKP rescues surface expression of these mutants.

### Gating properties of rescued mutant K_ATP_ channels

Some mutations that impair channel trafficking can also impact channel gating (32), it is therefore essential to verify that trafficking-deficient channels rescued to the cell surface maintain proper gating in response to ATP, MgADP, and diazoxide. Accordingly, we performed inside-out patch-clamp recordings on COSm6 cells transfected with mutant channels and treated overnight with AKP. We have previously shown that AKP inhibits channel activity in a readily reversible manner (22). To evaluate gating properties of AKP-rescued surface mutant channels, recordings were made after cells were incubated in AKP-free medium for at least 15 minutes to ensure removal of the drug and therefore its inhibitory effect on the channel. Following AKP overnight rescue and subsequent washout, K_ATP_ currents were detected in all six trafficking mutants, even for L171P and G228D which did not show obvious rescue by AKP in western blots. Possibly, western blots are not as sensitive for detecting low levels of rescue.

Representative current traces for N32K, P133R, and W143R mutants in response to ATP and MgADP or ATP and diazoxide are shown in Figure 6A and B. Quantification of the sensitivities to MgADP and diazoxide for all mutants, expressed as a percentage of currents relative to a nucleotide-free bath solution, is presented in Figure 6C. The data revealed that the MgADP and diazoxide sensitivities of the N32K, W143R, L171P, and Y230C mutants were comparable to those of WT channels. However, the P133R mutant showed a significant reduction in both MgADP and diazoxide sensitivities, and the G228D mutant exhibited a marked reduction in diazoxide response (Figure 6C). These findings indicate that while some trafficking mutants rescued to the cell surface retain the ability to respond to MgADP or diazoxide stimulation similarly to WT channels, others, such as P133R and G228D have compromised function.

**Figure 6.**
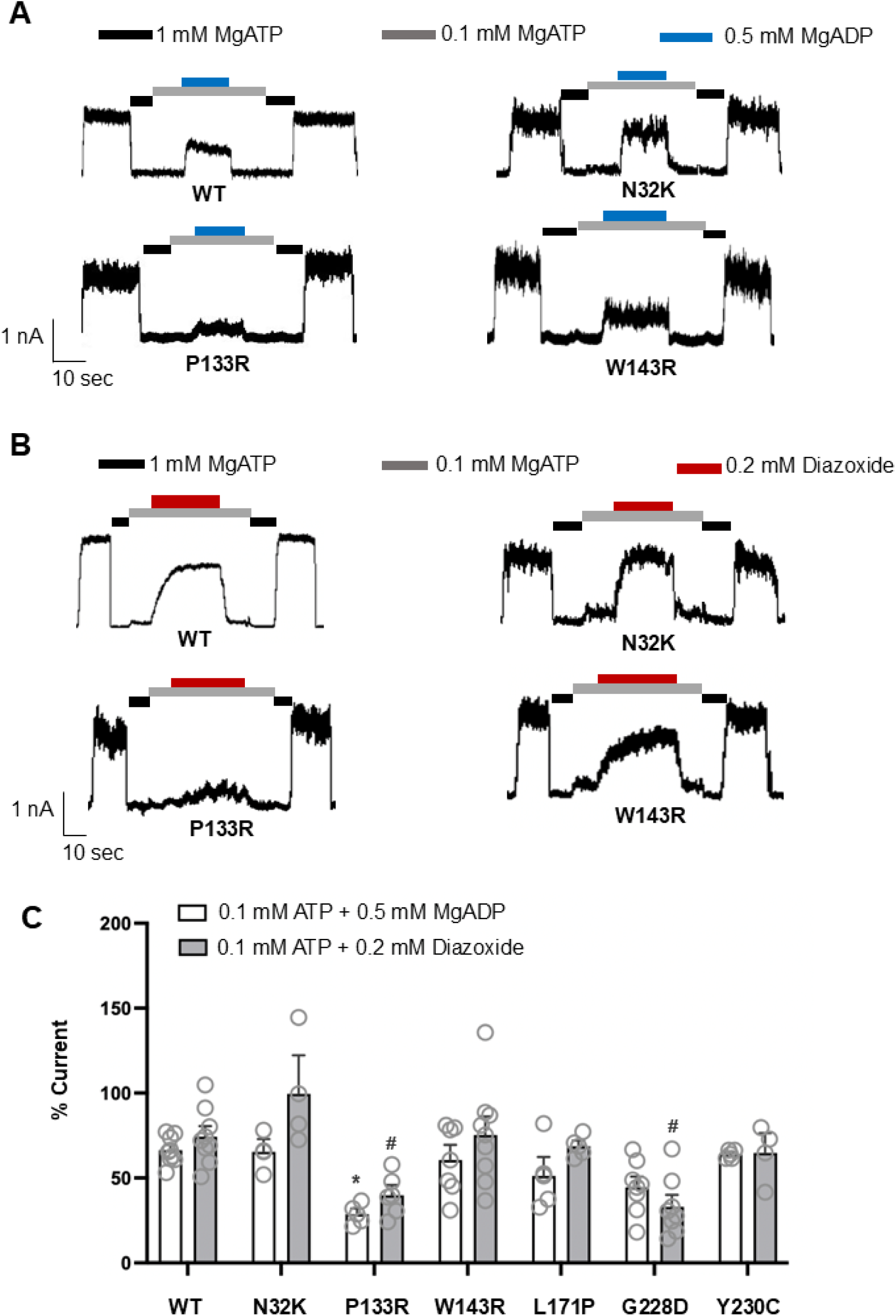
Gating properties of trafficking mutant channels rescued to the cell surface by Aekatperone assessed by inside-out patch-clamp recording. COSm6 cells expressing N32K-, P133R-, W143R-, L171P-, G228D-or Y230C-SUR1 mutant channels were incubated overnight with 100 μM Aekatperone (AKP) to rescue mutant channels to the cell surface. Before conducting the inside-out patch-clamp recordings, AKP was removed from the culture medium for at least 15 minutes, following the protocol described in “Experimental Procedures.” Cells expressing WT channels served as a control. (A) Representative current traces from WT, N32K, P133R, and W143R channels recorded in K-INT solution, with ATP and MgADP added as specified by the bars above the recordings. (B) Current traces from WT, N32K, P133R, and W143R channels recorded in K-INT solution, with or without ATP and diazoxide, as specified. For both panels (A and B), the recordings were conducted at a holding potential of −50 mV, with inward currents represented as upward deflections. (C) Quantification of the WT, N32K, P133R, W143R, L171P, G228D or Y230C channel response to 0.1 mM ATP + 0.5 mM MgADP or 0.1 mM ATP + 0.2 mM diazoxide. The currents were normalized to those measured in K-INT solution alone. Each bar represents the mean ± SEM (error bars) of at least five patches, with individual measurements indicated by grey circles (n). Significant difference (* or #, p < 0.05) was observed between the P133R and WT MgADP responses, and between P133R or G228D and WT Diazoxide responses as determined by a one-way ANOVA followed by Tukey’s multiple comparisons test.

### Assessing functional recovery of mutant K_ATP_ channels rescued to the cell surface in intact cells

The above results predict that following reversible pharmacochaperone rescue and washout, mutant channels, at least N32K, W143R, L171P, and Y230C, will respond to metabolic signals. To test this, we assessed channel activity in response to metabolic inhibition in intact cells by Rb^+^ efflux assays (23). COSm6 cells were transiently transfected with each of the six newly identified trafficking mutants: N32K, P133R, W143R, L171P, G228D, and Y230C. Following overnight treatment with 100 μM AKP the cells were subjected to Rb^+^ efflux assays, as described in Experimental procedures and in Figure 7A. AKP was washed out by incubating cells in AKP-free RbCl-containing medium for 20 minutes prior to the assay to ensure AKP unbinding and removal of any residual channel inhibition and excluded during the efflux assay. In parallel, cells treated overnight with 10 μM GBC or 200 μM Tolb and subjected to the same experimental procedure were included for comparison. All trafficking mutants exhibited significantly higher response to metabolic inhibition following treatment with AKP and Tolb, both reversible K_ATP_ inhibitor pharmacochaperones, although the magnitude of response varied across the mutations (Figure 7B). In contrast, cells treated with GBC, an irreversible K_ATP_ inhibitor pharmacochaperone, showed a reduced Rb^+^ efflux compared to vehicle-treated control cells, especially for channels that had significant activity in the absence of pharmacochaperones, including WT, N32K, P133R, and Y230C. This reduction is due to GBC inhibition of channels already present at the cell surface. These results emphasize the importance of using pharmacochaperones with reversible inhibitory effects for surface expression rescue and functional recovery of trafficking mutants.

**Figure 7.**
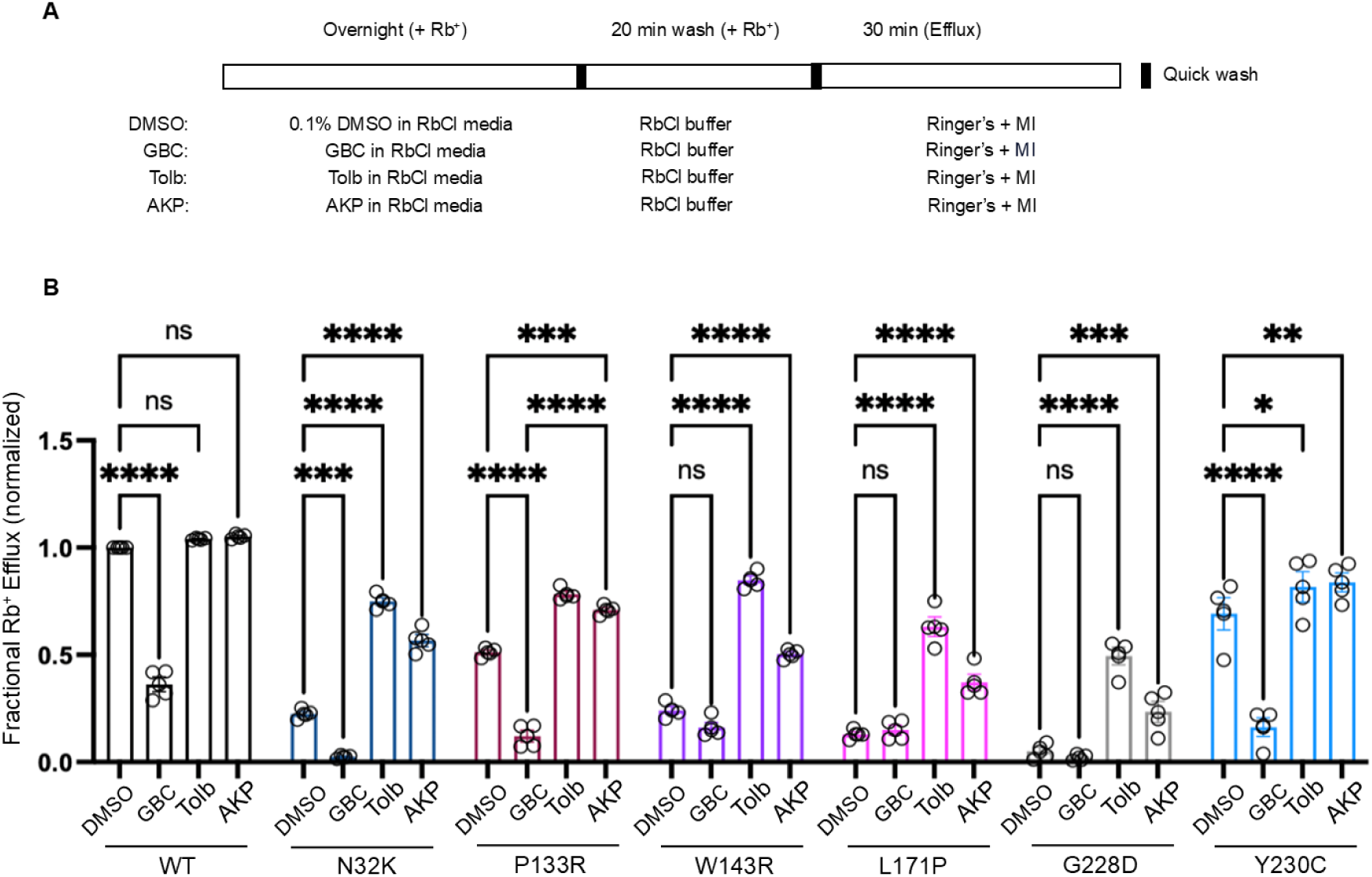
Functional rescue of trafficking mutant channels by various pharmacochaperones assessed by Rb⁺ efflux assays. (A) Schematic of the experimental protocol. COSm6 cells expressing WT or trafficking mutant K_ATP_ channels were treated with 0.1% DMSO (vehicle control), 10 μM glibenclamide (GBC), 200 μM tolbutamide (Tolb), or 100 μM Aekatperone (AKP) for 16 hours in the presence of Rb^+^. Prior to efflux measurements, cells were washed for 20 minutes in an RbCl-containing buffer that lacked any pharmacological compounds. Rb^+^ efflux was then measured over a 30-minute period in Ringer’s solution, either with or without metabolic inhibitors, MI (2.5 μg/ml oligomycin and 1 mM 2-deoxy-D-glucose). (B) Bar graph showing the fractional Rb^+^ efflux results of COSm6 cells that were transfected with WT or mutant (N32K, P133R, W143R, L171P, G228D, or Y230C) K_ATP_ channel plasmids. Untransfected cells were included to establish the background Rb⁺ efflux, which was subtracted from efflux in transfected cell groups. Bars represent the mean ± SEM from at least three independent biological replicates, with individual data points shown as circles. Data were normalized to the fractional Rb⁺ efflux of WT-expressing cells treated with metabolic inhibitors in the absence of pharmacochaperone treatment. Statistical significance was evaluated using one-way ANOVA in Prism, with α = (0.05). Significance levels are indicated as follows: \**p* < 0.05, \*\**p* < 0.01, \*\*\**p* < 0.001, \*\*\*\**p* < 0.0001.

We then assessed the diazoxide response of pharmacochaperone-rescued trafficking mutants using the Rb⁺ efflux assay using the same experimental design described above, but instead of adding metabolic inhibitors to the efflux solution, we added 200 μM diazoxide (see **Experimental procedures**; Figure 8A). Trafficking mutants which showed little SUR1 upper band in western blots (Figure 3A, B) had minimal diazoxide response in vehicle-treated cells as expected, in contrast to WT channels (Figure 8B). Some diazoxide response was seen for P133R and Y230C even in vehicle-treated cells, likely due to partial surface expression, as shown by western blot where the upper band was present, though at lower levels than WT (Figure 3A). Tolb and AKP overnight treatment followed by washout increased diazoxide response, with the extent of enhancement across the different mutations mirroring that seen for metabolic inhibition (see Figure 7), although for L171P and G228D, the AKP rescue did not result a statistically significant increase in diazoxide response. Together, these results further extend the effectiveness of reversible K_ATP_ inhibitor pharmacochaperone such as Tolb and AKP in enhancing metabolic inhibition and diazoxide response to the new K_ATP_ trafficking mutants identified in this study.

**Figure 8.**
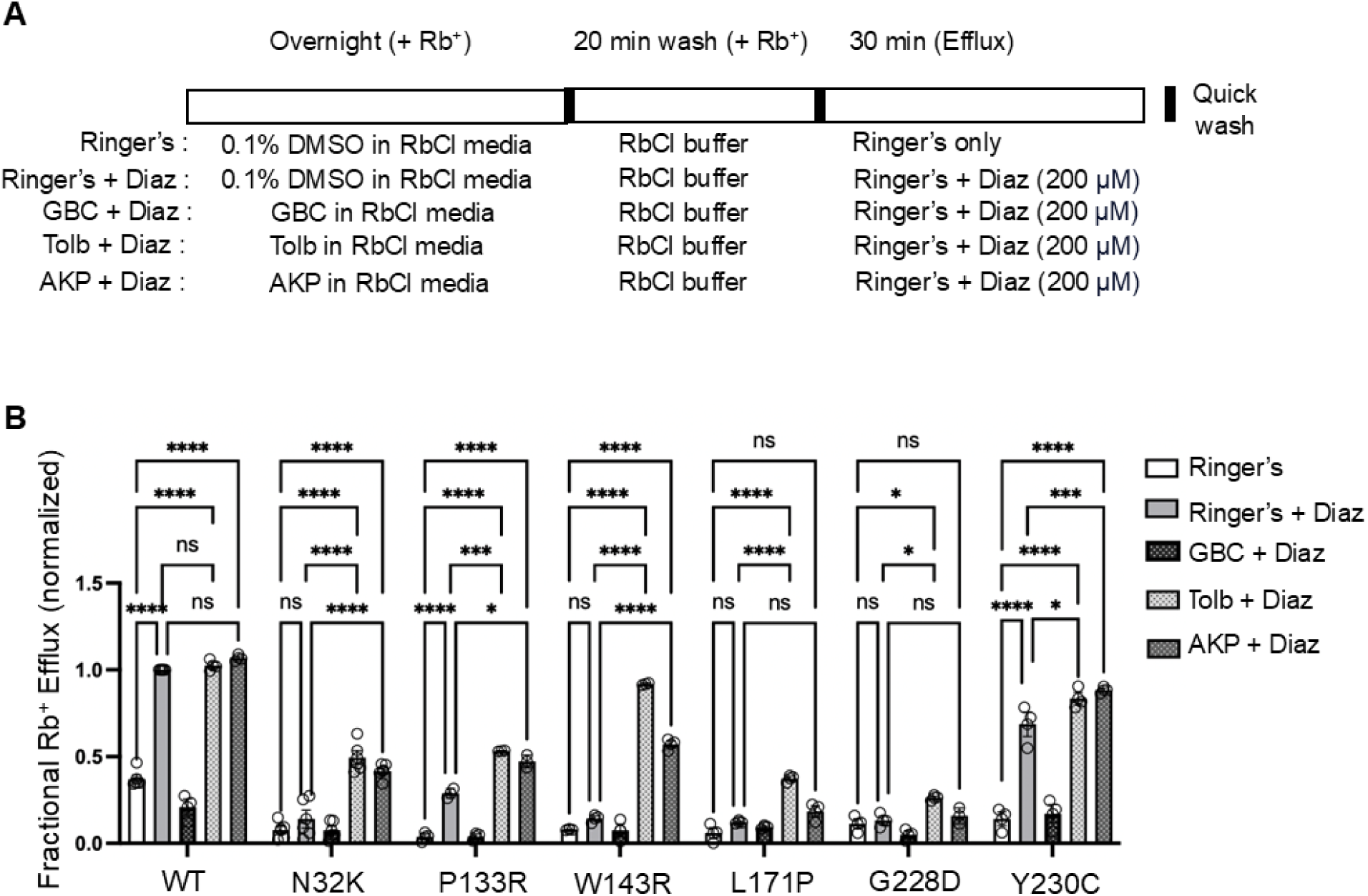
Enhanced acute diazoxide response following overnight pharmacochaperone treatment in COSm6 cells expressing trafficking mutants. (A) Schematic of the experimental protocol. COSm6 cells expressing WT or trafficking mutant K_ATP_ channels were treated with 0.1% DMSO (vehicle control), 10 μM glibenclamide (GBC), 200 μM tolbutamide (Tolb), or 100 μM Aekatperone (AKP) for 16 hours in the presence of Rb^+^. Prior to efflux measurements, cells were washed for 20 minutes in an RbCl-containing buffer that lacked any pharmacological compounds. Rb^+^ efflux was then measured over a 30-minute period in Ringer’s solution, either with or without 200 μM diazoxide (Diaz). Note that diazoxide was added to the Ringer’s solution during the efflux assay but was not present during the overnight incubation. (B) Bars represent the mean ± SEM from at least three independent biological replicates, with individual data points shown as circles. Statistical significance was determined using one-way ANOVA (α = 0.05). Significance levels are indicated as follows: *p < 0.05, **p < 0.01, ***p < 0.001, ****p < 0.0001.

## Discussion

K_ATP_ channel mutations represent the most frequent genetic cause of HI (7, 33). Unlike cystic fibrosis, which is predominantly caused by a single recessive misfolding mutation ΔF508 in CFTR (34, 35), K_ATP_-HI exhibits significant genetic heterogeneity, with numerous mutations distributed throughout the K_ATP_ channel subunits impacting channel expression, gating, or both (7–9, 36). The diverse genetic variants in K_ATP_-HI and their complex clinical manifestations— ranging from dominant or recessive inheritance, focal or diffuse disease, to compound mutations or benign polymorphisms—make their pathogenic classification challenging, which hinders rapid diagnosis and mechanism-based treatment. The spectrum of HI-associated mutations in K_ATP_ channels remains incompletely defined. Indeed, approximately 70% of missense mutations identified to date represent new variants of unknown significance (8, 9), emphasizing the need for ongoing discovery and characterization of new mutations. Our study presented here provides detailed biochemical and functional data on a cohort of newly identified SUR1 missense variants identified in seven diazoxide-unresponsive HI patients presenting with focal or diffuse disease and carrying single or compound heterozygous mutations. The data validate the loss-of-function effects of these variants thereby establishing their pathogenic role in HI and clarify the relative contributions of compound mutations in disease phenotype and diazoxide response. Moreover, the study highlights the critical role of SUR1-TMD0/L0 in channel trafficking and gating, and it identifies candidate mutations in this domain for potential pharmacochaperone therapy.

### Genotype-phenotype correlation in patients with compound heterozygous mutations

Genotype-phenotype correlation is important for accurate diagnosis, treatment, and genetic counseling. Of the seven diazoxide-unresponsive patients, three have focal disease with single mutations that are clearly pathogenic based on our biochemical and functional studies. However, four have diffuse disease and carry compound heterozygous mutations on separate alleles. One patient has a paternal N32K mutation and a maternal truncation mutation Q954X, and a second patient has a maternal G228D mutation and a paternal frameshift mutation, D1193Mfs. Q954X and D1193Mfs are predicted to be nonfunctional, therefore the disease phenotype is likely determined by N32K or G228D, both of which show severe trafficking defects. The other two patients, one harbors a non-maternal Y124F mutation with a maternal C1491Y mutation, and the other has a maternal Y230C mutation and a paternal G1384R mutation. Our study shows that while both Y124F and Y230C reduced channel response to MI and diazoxide in Rb efflux assays (Figure 2A), they retained residual MI and diazoxide response (> or ∼50% of WT levels), which would predict mild disease and diazoxide responsiveness in the patients (37). In contrast, both G1384R and C1491Y showed almost no or little response to MI and diazoxide, and in western blots showed no SUR1 upper band indicating severe trafficking defects, which are consistent with the clinical phenotype observed in patients. The dominant roles of C1491Y and G1384R over Y124F and Y230C both in SUR1 maturation and K_ATP_ function are further demonstrated by Y124Y/C1491Y or Y230C/G1384R co-expression studies (Figures 2, 3). These studies suggest C1491Y- and G1384R-SUR1 can co-assemble with Y124F- and Y230C-SUR1 respectively to exert their dominant-negative effects on channel trafficking and function.

### SUR1-TMD0/L0 in K_ATP_ channel trafficking and gating

In the K_ATP_ channel complex, the SUR1-TMD0/L0 domain interfaces directly with Kir6.2 and is critical for channel assembly, thereby trafficking to the cell surface. In addition, the domain is involved in K_ATP_ channel gating by ATP and PIP_2_, which inhibits and stimulates the channel respectively. Specifically, L0 contributes to inhibitory ATP binding at Kir6.2 (38–41), while TMD0 participates in PIP_2_ binding to stabilize Kir6.2 in an open conformation (42). Moreover, conformational remodeling at the TMD0/L0 and Kir6.2 interface has been observed comparing inhibitory ATP-bound closed channel structure and the structure of open channels where SUR1-NBDs are bound to MgADP/MgATP and dimerized (43, 44). In previous studies we have identified many TMD0 mutations which impair SUR1 maturation and K_ATP_ trafficking to the plasma membrane and which can be rescued to the cell surface by pharmacochaperones (17, 19, 29, 30). Most of these trafficking mutants rescued to the cell surface showed normal gating properties like WT, but some showed altered gating (17).

In the current study, all seven newly identified SUR1-TMD0/L0 mutations were found to reduce the mature SUR1 band on western blots, although the mild reduction in Y124F (20±13%) did not reach statistical significance. Y124F did, however, show a significant reduction in MgADP response but no obvious change in sensitivity to ATP inhibition, therefore the reduced MgADP response is likely the primary explanation for its reduced activity in Rb^+^ efflux assays under metabolic inhibition condition. Among the trafficking mutants, P133R and G228D rescued to the cell surface also showed altered gating. While sensitivity to ATP inhibition appeared unchanged by either mutation, both P133R and G228D reduced MgADP and diazoxide response, although the reduction in MgADP response for G228D did not reach statistical significance (*p* = 0.1552; Figure 6C). Both MgADP and diazoxide stimulate K_ATP_ channels by binding to the ABC-core structure of SUR1; the former binds to NBD2 and promotes NBD dimerization, and the latter binds to the transmembrane domains of the SUR1-ABC core and stabilizes SUR1 in an NBD dimerized conformation (16). Since Y124F, P133R, and G228D reside in TMD0/L0 outside the SUR1-ABC core (see Figure 1), they likely reduce MgADP and diazoxide responses by interfering with the structural changes that couple SUR1-NBD dimerization with Kir6.2 opening.

Alternatively, they may interfere with PIP_2_ binding or gating, which ultimately opens the channel, to indirectly reduce channel response to MgADP or diazoxide. Our findings reinforce the importance of SUR1-TMD0/L0 in channel assembly and trafficking, and they provide functional evidence to support structural observations that suggest SUR1-TMD0/L0 couples ligand-induced structural changes at the SUR1-ABC core to Kir6.2 gating.

### Pharmacochaperone rescue of trafficking mutants: therapeutic implications

K_ATP_ trafficking mutations are common in HI (7, 17), yet there is currently no clinically approved pharmacotherapy for patients affected by such mutations. Our previous demonstration that K_ATP_ channel inhibitors can act as pharmacochaperones and improve surface expression of trafficking mutants located in SUR1-TMD0 offers a potential solution, albeit a partial one, as not all TMD0 trafficking mutations are efficiently rescued and trafficking mutations in the large SUR1-ABC core are not rescued by these inhibitors. For this reason, it is important to identify trafficking mutations that are amenable to pharmacochaperone rescue. Once K_ATP_ trafficking mutants are rescued to the cell surface by pharmacochaperones, the inhibitor chaperones need to be removed to recover channel function for therapeutic applications. This requires that the inhibitory effects of the pharmacochaperone be rapidly reversible. Early work found that the low affinity sulfonylurea Tolb and the anti-convulsant carbamazepine have such desired properties (19, 21). However, Tolb is off the market due to concerns of its cardiotoxicity possibly through off-target effects on hERG channels (45), while carbamazepine has many other non-K_ATP_ targets (46). Using structure-based virtual screening combined with functional validation we have recently discovered a new reversible K_ATP_ inhibitor pharmacochaperone, AKP, which in our initial study was shown to rescue the surface expression and function of several know TMD0 trafficking mutants (22).

In this study, we further tested the effects of AKP on newly identified trafficking mutations. We found that while four of the six TMD0/L0 trafficking mutations including N32K, P133R, W143R, and Y230C showed clear response to AKP and to Tolb and GBC, the other two, L171P and G228D, showed no obvious increase in the SUR1 upper mature band. Further, the two trafficking mutations in the SUR1-ABC core, G1384R and C1491Y, did not show an increase in SUR1 upper band. This pattern is consistent with published studies (17, 19, 29, 30) and steer the focus on pharmacochaperone therapy to SUR1-TMD0/L0 trafficking mutations. In Rb^+^ efflux assays, the reversible inhibitor pharmacochaperones AKP and Tolb showed differential efficacies in restoring TMD0/L0 trafficking mutant response to MI and diazoxide. At the concentrations tested, AKP is as effective as Tolb for N32K, P133R, and Y230C, but AKP is less effective than Tolb for W143R, L171P, and G228D. Worth noting, the extent of functional rescue does not always match the extent of upper SUR1 band rescue in western blots. This could be explained by the interplay between channel surface expression and gating properties. For example, the rescued P133R mutant channels showed reduced MgADP and diazoxide response and G228D mutant channels showed reduced diazoxide response in electrophysiology experiments. The impaired gating would reduce the function of mutant channels rescued to the cell surface. We also found that although L171P and G228D showed no clear increase in the upper SUR1 band in western blots upon pharmacochaperone treatments, they did show some functional recovery in Rb efflux assays, which likely reflects the different sensitivities of the methods used to monitor pharmacochaperone effects.

Our study expands the HI-causing SUR1 trafficking mutations amenable to surface expression and functional rescue by AKP, providing the impetus to further test the safety and efficacy of AKP in relevant cell and animal disease models.

### Summary

Timely diagnosis and treatment of HI are crucial for preventing life-threatening hypoglycemia and its associated complications. Our study highlights the pathogenic diversity of K_ATP_ channel mutations and the value of detailed characterization of disease associated genetic variants in guiding personalized therapeutic approach. Pharmacochaperone therapy presents a promising strategy for patients with K_ATP_ channel trafficking defects who do not respond to conventional diazoxide therapy. More research to further expand our knowledge base of trafficking mutations amenable to pharmacochaperone rescue, and to optimize the efficacy and minimize off-target effects of K_ATP_ pharmacochaperones will aid in the translation of this therapeutic concept to clinical applications.

## Experimental procedures

### Genetic and clinical studies

The subjects in this study were patients referred to the Children’s Hospital of Philadelphia (CHOP) Congenital Hyperinsulinism Center, as well as patients reported in the literature (see **Table 1**). CHI patients were classified as diazoxide-unresponsive if the cardinal feature of HI, hypoketotic hypoglycemia, could not be reversed by diazoxide at a dosage of 15 mg/kg/day (for at least 5 days) (4). Most of these patients ultimately required surgical pancreatectomy. Clinical information was obtained from their medical records. Written informed consent was secured from the parents of all probands included in the study. The study protocol was reviewed and approved by the Institutional Review Board of CHOP.

Mutation analysis was performed either in commercial laboratories or as part of a research study at the Children’s Hospital of Philadelphia. Peripheral blood samples were collected from patients to isolate genomic DNA (5 PRIME, Gaithersburg, MD). Coding regions and intron/exon splice junctions were amplified and sequenced directly using an ABI 3730 capillary DNA analyzer (Applied Biosystems, Carlsbad, CA). *ABCC8* nucleotide positions and corresponding SUR1 amino acid numbers are based on the sequence provided by Nestorowicz et al. (47), which includes the alternatively spliced exon 17 sequence (NCBI accession number L78224).

### Molecular biology

In all experiments, human FLAG epitope-tagged SUR1 (referred to as f-SUR1) cDNA in pCMV6b (generously provided by Dr. Joseph Bryan) and human Kir6.2 in pcDNA3.1 were used. Point mutations were introduced using the QuikChange site-directed mutagenesis kit (Stratagene). The FLAG epitope (DYKDDDDK) was inserted at the N terminus of hamster SUR1 cDNA, as described previously (31). Prior studies have confirmed that placing the FLAG epitope at the extracellular N terminus of SUR1 does not impact channel assembly or function (14, 30). All mutations were verified by DNA sequencing, and mutant clones from two independent PCR reactions were analyzed in each experiment to prevent artifacts from undesired PCR-introduced mutations.

### Transfection of COSm6 cells with recombinant DNA

COSm6 cells were plated in 6-well tissue culture plates at approximately 70% confluency and transfected with 1.2 μg of wild-type (WT) or mutant SUR1, along with 1.2 μg of WT Kir6.2 per well, using FuGENE 6 (Promega) according to the manufacturer’s protocol. To simulate heterozygous expression, in some experiments, COSm6 cells were co-transfected with mutant SUR1 cDNAs (e.g., Y124F/C1491Y or Y230C/G1384R) in a 1:1 ratio, maintaining a total of 1.2 μg of SUR1 DNA, alongside 1.2 μg of WT Kir6.2. Importantly, plasmids were first pre-mixed in Opti-MEM in a separate tube before the addition of FuGENE® to ensure efficient uptake of multiple plasmids required for proper K_ATP_ channel expression.

### Immunoblotting

COSm6 cells were transfected with SUR1 and Kir6.2 using FuGENE^®^6 reagent. After 48 hours, cells were lysed in a buffer containing 50 mM Tris·HCl (pH 7.0), 150 mM NaCl, and 1% Triton X-100, supplemented with CompleteTR protease inhibitors (Roche Applied Science), on ice for 30 minutes. Proteins from the cell lysates were then separated by SDS-PAGE on a 7.5% gel, transferred onto a nitrocellulose membrane, and probed with a rabbit anti-SUR1 antibody, which was raised against a C-terminal peptide of SUR1 (KDSVFASFVRADK) (30). This was followed by incubation with horseradish peroxidase (HRP)-conjugated secondary antibodies (Amersham Biosciences). Detection was carried out using enhanced chemiluminescence (Super Signal West Femto, Pierce). Tubulin was used as a loading control and was also probed in the blots.

### Immunofluorescence staining

COSm6 cells were cultured on coverslips and transfected with human f-SUR1 and human Kir6.2 constructs. Cells were treated for 16 hours with either 0.1% DMSO, 10 μM Glibenclamide (GBC), 200 μM Tolbutamide (Tolb), or 100 μM Aekatperone (AKP) before proceeding with immunofluorescence staining. To label channels present on the cell surface, cells were washed with ice-cold phosphate-buffered saline (PBS) containing 137 mM NaCl, 2.7 mM KCl, 10 mM Na₂HPO₄, and 1.8 mM KH₂PO₄ at pH 7.4. They were then incubated with M2 anti-FLAG antibody (10 μg/ml in Opti-MEM with 0.1% BSA) for 1 hour at 4°C. Following this, cells were washed with ice-cold PBS, fixed in 4% paraformaldehyde for 10 minutes on ice, and washed three times with cold PBS. The fixed cells were then blocked for 1 hour in PBS containing 2% BSA and 1% normal goat serum, followed by a 1-hour incubation with Alexa Fluor 546-conjugated goat anti-mouse secondary antibody (Invitrogen; diluted 1:300 in blocking buffer) at room temperature. For staining total cellular FLAG-SUR1, cells were fixed with 4% paraformaldehyde on ice for 10 minutes before incubating with the anti-FLAG antibody, followed by Alexa Fluor 546-conjugated goat anti-mouse secondary antibody. Cells were subsequently washed twice with PBS, and the coverslips were mounted onto microscope slides using Vectashield Mounting Medium for Fluorescence with DAPI. Imaging was performed using an Olympus confocal microscope.

### Non-radioactive Rb^+^ efflux assays

COSm6 cells were transiently transfected with various combinations of human wild-type (WT) or mutant SUR1 and Kir6.2 cDNAs, while untransfected cells served as background controls. Cells were cultured overnight in medium containing 5.4 mM RbCl. The following day, cells were washed twice with PBS without RbCl.

For experiments assessing the effect of mutations on K_ATP_ channel function (Figures 2), Rb⁺ efflux was measured by incubating cells in Ringer’s solution (5.4 mM KCl, 150 mM NaCl, 1 mM MgCl₂, 0.8 mM NaH₂PO₄, 2 mM CaCl₂, 25 mM HEPES, pH 7.2) either under baseline conditions (only Ringer’s solution), metabolic inhibition, (2.5 μg/ml oligomycin and 1 mM 2-deoxy-D-glucose) (Ringer’s + MI), or in the presence of 200 μM diazoxide (Ringer’s + Diazoxide), for 30 minutes at 37°C. For experiments evaluating the rescue of channel function at the cell surface after pharmacochaperone treatment with drugs like glibenclamide, tolbutamide, and Aekatperone (Figure 7), the drugs were added to the RbCl-containing medium overnight (16 h). An additional wash step with RbCl wash buffer (5.4 mM RbCl, 150 mM NaCl, 1 mM MgCl₂, 0.8 mM NaH₂PO₄, 2 mM CaCl₂, 25 mM HEPES, pH 7.2) at 37°C for 20 minutes was included to remove the drugs prior to the efflux assay. Rb⁺ efflux was then measured by incubating the cells in Ringer’s solution with only metabolic inhibitors, without Aekatperone (22, 23). Similarly, for experiments assessing the role of pharmacochaperones in enhancing the response to diazoxide (Figure 8), drugs were included in the RbCl-containing medium overnight, followed by a drug washout step. The efflux assay was then performed in Ringer’s solution containing 200 μM diazoxide. For each experiment, technical duplicates were included, and the average value was used as the experimental result. At least three separate transfections were performed for each experimental condition as biological repeats, as detailed in the figure legends (number of biological repeats represented as small circles on the graphs). Data are presented as mean ± SEM.

### Patch-clamp recordings

COSm6 cells were transfected using FuGENE^®^6 and plated onto coverslips. To identify transfected cells, GFP cDNA was co-transfected along with SUR1 and Kir6.2. Patch-clamp recordings were conducted 48–72 hours post-transfection, and all experiments were performed at room temperature as previously described (30). Micropipettes were pulled from non-heparinized Kimble glass (Fisher) using a horizontal puller (Sutter Instrument Co., Novato, CA), with electrode resistance typically between 1–2 megaohms when filled with K-INT solution (composition below). Inside-out patches were voltage-clamped using an Axopatch 1D amplifier (Axon Inc., Foster City, CA).

The standard bath (intracellular) and pipette (extracellular) solution, referred to as K-INT, contained 140 mM KCl, 10 mM K-HEPES, 1 mM K-EDTA, pH 7.3. ATP and ADP were added as potassium salts. Currents were recorded at a membrane potential of −50 mV (pipette voltage set at +50 mV). Data acquisition and analysis were carried out using pCLAMP10 software (Axon Instruments) and Microsoft Excel, with results presented as means ± S.E.M. For experiments assessing the ATP inhibitory responses of the F44V mutation, ATP response was measured in a Mg²⁺-free solution with EDTA included. In contrast, for experiments evaluating the gating properties of the channel, MgCl₂ was added to the ATP, ADP, or diazoxide-containing solutions, maintaining a free Mg²⁺ concentration of approximately 1 mM (48) .

### Statistics

Data are presented as mean ± SEM. Statistical comparisons between two groups were performed using Student’s t-test, while comparisons among three or more groups were analyzed with one-way ANOVA, followed by Tukey’s or Dunnett’s post-hoc tests for multiple comparisons, as indicated in the figure legends. Analyses were conducted using GraphPad Prism 10, and differences were considered statistically significant if *p* ≤ 0.05.

## Data availability

All data are contained within the manuscript.

## Acknowledgements

We thank Drs. Camden Driggers and Bruce Patton for helpful discussions.

## Author contributions

AE conceived the project, designed, and performed experiments, analyzed data, prepared all figures, and wrote the manuscript; YYK performed electrophysiological recordings, KB, CS and DDL performed genetic analysis for patients and provided us with all patient information, ZY performed experiments; SLS conceived the project, analyzed data, wrote, and edited the manuscript.

## Funding and additional information

The work is supported by the National Institutes of Health grant R01GM145784 (to SLS) and R01DK056268 (to DDDL). AE is recipient of the Egyptian government predoctoral scholarship GM 1109 and YYK is recipient of the American Heart Association Postdoctoral Fellowship Award 25POST1373218.

The content of the manuscript is solely the responsibility of the authors and does not necessarily represent the official views of the National Institutes of Health.

## Abbreviations

K_ATP_: ATP-sensitive potassium channel
HI: congenital hyperinsulinism
cryoEM: cryo-electron microscopy
Kir6.2: inward-rectifying potassium channel 6.2
SUR1: sulfonylurea receptor 1
ATP: adenosine triphosphate
ADP: adenosine diphosphate
GBC: glibenclamide
Tolb: tolbutamide
AKP: Aekatperone
ABC protein: ATP-binding cassette protein
NBD: nucleotide binding domain of SUR1
TMD0: transmembrane domain 0
ER: endoplasmic reticulum
CFTR: cystic fibrosis transmembrane conductance regulator protein

